# The distribution of clade size in a coalescent diversification model

**DOI:** 10.1101/2024.11.03.621532

**Authors:** Yexuan Song, Caroline Colijn, Ailene MacPherson

## Abstract

Characterizing the patterns and determinants of biological diversity is a central aim of evolutionary biology. Doing so requires developing expectations for clade size, the relationship between the number of species in a clade and its age, for standard diversification models, such as the coalescent or birth-death process. These expectations are necessary for identifying diversity outliers, specious or depauperate clades in the macroevolutionary context or transmission clusters of particular or little public health concern in the epidemiological context, and for testing alternative diversification hypotheses. Here, we derive a closed-form expression for the distribution of clade sizes under the widely used Kingman coalescent diversification model and extend the results to allow the number of niches (the effective population size) to vary through time. This result complements analogous results for the birth-death model in that it provides expectations for the joint distribution of clade sizes for a case where diversification is strongly density-dependent.

## 1. Introduction

“How large should a clade be?” is a longstanding and enduring question in evolutionary biology [1, 2, 3] with wide-ranging applications in paleobiology [4], epidemiology [5], conservation [6], and cellular biology [7]. Developing expectations for clade size under standard diversification models such as the Yule [1], or the birth-death model [2, 3] is necessary, for example, to identify diversity outliers (e.g., specious or depauperate clades) and to test and select between alternative diversification hypotheses. In the macroevolutionary context, diversity outliers include clades shaped by adaptive radiations and those inhabited by ‘living fossils.’ Classic examples of the former include Anolis lizards [8] and Cichlid fishes [9]. In contrast, examples of the latter include tailed frogs (*Ascaphus truei*) [10] and Gars (*Lepisosteidae*) [11]. In an epidemiological context where the diversity of viral lineages is considered, large clades may indicate transmission clusters of particular concern and may provide insights on identifying high-risk populations [12, 13]. As such, theoretical expectations are needed to identify outlier clusters and how the distribution of cluster size depends on the epidemiological dynamics (e.g., a growing epidemic vs. a constant endemic prevalence). Clade sizes can also form the bases of tests of alternative diversification hypotheses in macroevolution. Examples include testing whether diversification is “bounded” due to competition for a finite niche space or “unbounded” [14, 15].

Beginning with Yule [1], models have long been used to quantify diversity [3]. Many biodiversity questions are naturally framed regarding tree or clade size, the relationship between the number of lineages in a tree/clade and its height. For example, Yule was interested in the expected sizes of genera, using his speciation-only model to propose that after time *T*, the number of species should follow a geometric distribution, a relationship later termed the “hollow curve” [1]. Despite the initial focus of modelling efforts (e.g.[1, 16]), clade size is rarely considered in contemporary phylodynamics. Rather, methods for inferring diversification dynamics typically use branch lengths (speciation, extinction, and sampling times) [17, 18, 7, 19, 20] or tree imbalance [5, 21] to test alternative diversification hypotheses. Our recent work [3] considering clade size in the case of the birth-death model, however, demonstrated that the size of a phylogenetic tree may be particularly useful for assessing emergent biodiversity phenomena such as whether diversification is density-(in)dependent. Here we develop expectations for the size of a single clade or a combination of clades under the coalescent diversification model.

The Kingman Coalescent [22] is a widely used diversification model, particularly in epidemiology where its analytical tractability can facilitate the rapid inference of dynamics of disease spread [23, 24, 25]. Derived as a large-population size limit of the Wright-Fisher model, the Kingman Coalescent describes the genealogical relationships between a set of sampled sequences. Although as a diversification model the coalescent no longer has an explicit evolutionary description as it does as a model of genealogies, it can be used as an approximation of diversification dynamics with strong density dependence. Specifically, it can be interpreted as a model of diversification where there is a large constant (or deterministically changing) number of filled ecological niches from which a small number of lineages are sampled. Speciation and extinction occur as species in one niche replace species in other niches. A summary of the classic diversification models can be found in Table D.3.

Here, we define the size of a clade at a ‘cut-off time’ *T* (red horizontal line in Fig. 1) as the size of the subset of the observed lineages at the present day which share a particular common ancestor at the cut-off time. As a result of the strong implicit density dependence in the diversification process in combination with the fixed total number of observed lineages, the size of two or more clades in the coalescent model will not be independent. As such, here, we derive both the (marginal) probability of observing a single clade of a specific size and the joint probability of observing two or more clades of specific sizes. This examination of clade size complements existing measures of diversity in a coalescent model that is grounded in population genetics; namely, Ewens’s sampling formula and the Site Frequency Spectrum (SFS) [26, 27, 28]. Ewens’s sampling formula and the SFS exemplify the use of marginal and joint probability distributions in quantifying diversity under the coalescent model. Both metrics consider the pattern of genetic diversity arising from mutations along the branches of a genealogy when mutations occur according to an infinite-alleles and infinite-sites model, respectively (see Fig. 1). A complete description of the (unfolded) SFS, for example, is summarized by the joint probability of observing *ζ*_1_singleton mutations (mutations present in a single observed lineage) and *ζ*_2_ doubletons etc., up to *ζ*_*n*_ mutations present in all *n* sampled lineages. For practical purposes, however, this joint distribution is often summarized by corresponding marginal distributions (e.g., the distribution of singletons).

**Figure 1:**
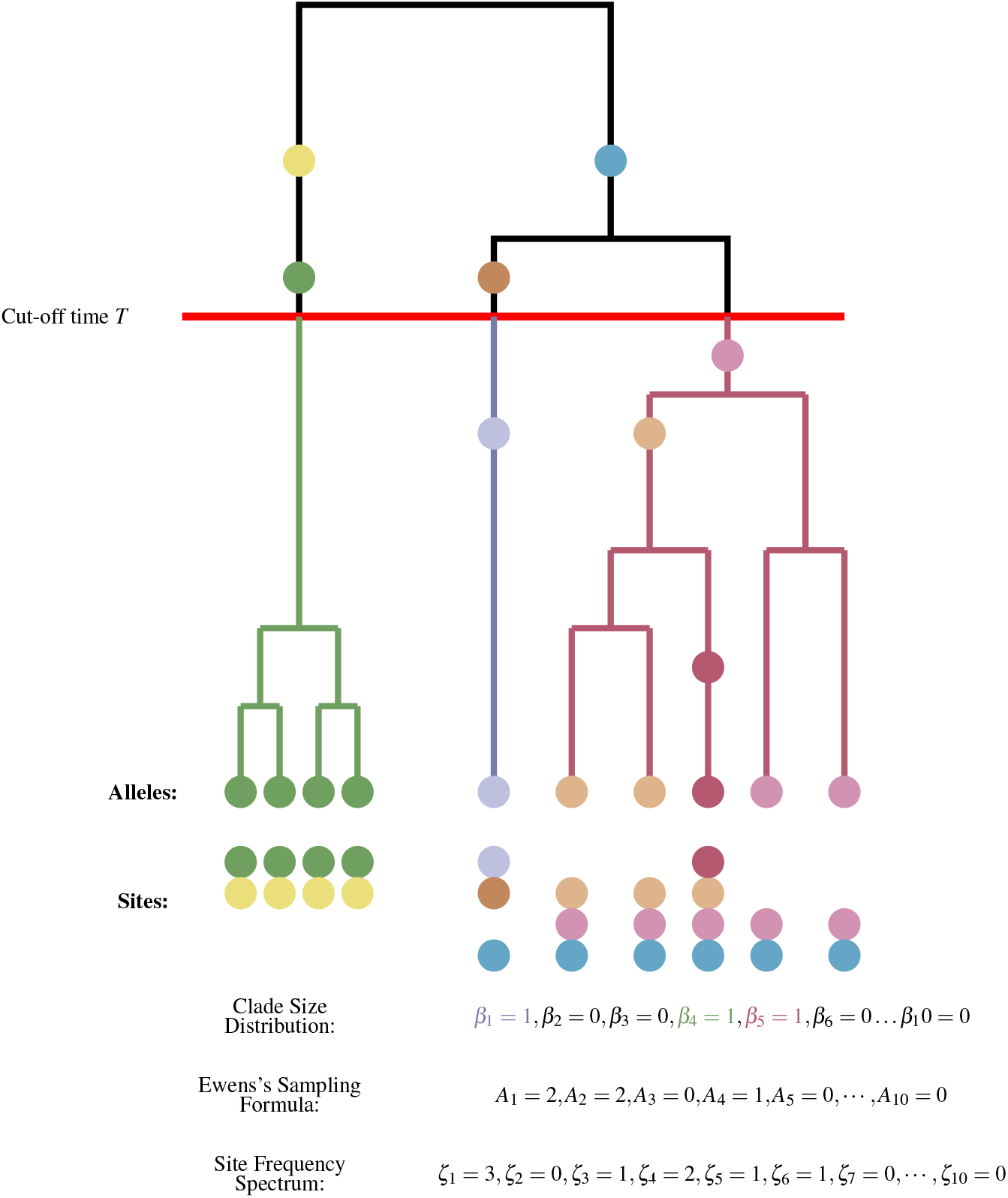
Comparing different metrics (Ewens’s Sampling formula, Site Frequency Spectrum, and Clade Size Distribution) of diversity in the coalescent model. Mutations (filled circles) occur along the edges of an genealogy with *n* = 10 observed lineages whereas coloured edges represent clades. Ewens’s sampling formula gives the joint probability, Pr(*A*_1_, *A*_2_, … *A*_*n*_) of observing *A*_1_ alleles once in the sample, *A*_2_ alleles represented twice and so on such that ∑_*i*_ *iA*_*i*_ = *n*. The SFS gives the joint probability, Pr(*ζ*_1_, *ζ*_2_, … *ζ*_*n*_), of observing *ζ*_*i*_ sites with the mutation observed *i* times in the sample. Finally, the clade size distribution gives the probability Pr(*β*_1_, *β*_2_, … *β*_*n*_ | *T*) of observing *β*_1_ clades of size 1, *β*_2_ clades of size 2, etc. given some cutoff time *T*.

In the next section, we derive analytical expressions for the marginal and joint distribution of clade sizes. To facilitate the application to a range of biological systems, we consider five different cases. We begin by deriving expectations for the marginal (Cases 1 and 2) and joint (Cases 3 and 4) distribution of clade sizes when the niche size *N* remains constant (Cases 1-4). We consider two different sampling schemes, one in which whole clades are sampled randomly (Cases 1 and 3) and one in which lineages (within clades) are chosen randomly (Cases 2 and 4). We then extend these results (Case 5), allowing the number of niches, hereafter referred to as the niche size, to vary through time, for example, as used in coalescent-based phylodynamic analyses [29]. The derivation of these joint distributions is particularly novel as there are no known analogous expressions in the case of the birth-death model when diversification is density-dependent. Hence, this coalescent expectation provides a valuable reference for evaluating biodiversity patterns in contexts such as epidemiology, where diversification is expected to be strongly density-dependent. We validate our analytical expressions using simulations and use the expressions to make predictions for biodiversity patterns. We conclude with a discussion of the implications of these distributions for applications in macroevolution and epidemiology.

## 2. Methods

Here we consider the evolutionary history of a small number of *n* lineages observed at the present-day evolving according to a neutral Wright-Fisher process. The number of niches, hereafter referred to as the niche size, at time *τ* before the present day is given by *N*(*τ*) where time is measured in units of *N*_ref_ generations (see Table 1). Given our focus on the coalescent as a tree prior, throughout we will primarily use ‘niches’, ‘lineages’, and ‘clades’ to refer to the features of the coalescent model. Alternative terminologies for the coalescent as a genealogy, in terms of integer partitions, and in the context of the Pólya Urn model are described in Table D.4 and used when necessary. We will initially assume (Cases 1-4) that the number of niches, *N*(*τ*) = *N*_ref_, is constant over time, an assumption we relax (Case 5) to allow for arbitrarily varying sizes. When the niche size is constant, *o* represents the total niche size. When niche size varies through time, *N*_ref_ represents a ‘reference niche size’. Assuming that *N*_ref_ ≫*n*, the ancestry of the observed lineages is then described by the classic Kingman Coalescent [22]. We then consider the clades formed by subsets of the *n* samples that share a common ancestor at a ‘cut-off time’ *T*, where *T* is measured backward in time in coalescent time units (see Fig. 1).

**Table 1:** List of notation.

| Variable | Description |
| --- | --- |
| $N(\tau)$ | The number of filled niches (gene copies) at time $\tau$ before the present day in coalescent units. $N_{\text{ref}}$ is used to refer to the constant or reference niche size. We refer to the number of filled niches as the “niche size”. |
| $n$ | The number of “observed lineages” (sequences) in the observed tree at the present day, $\tau = 0$. Throughout, we refer to these lineages as “observed” to distinguish them from the “sampling” of clades or “sub-sampling” of lineages within clades. |
| $T$ | The “cut-off time” measured in units of $N_{\text{ref}}$ generations at which clades are assessed. |
| $m$ | The size of a single “sampled” clade. |
| $C = (m_1, \dots, m_c)$ | A tuple giving the sizes of $c$ sampled clades where $m_j$ is the size of the $j^{\text{th}}$ clade ($1 \leq j \leq c$). Throughout, we use $c$ to denote the number of clades sampled where $1 \leq c \leq k$ where $k$ is the number of ancestors at the cut-off time. |
| $x = \{s_1, \dots, s_k\}$ | A partition of the integer $n$ of size $k$. Throughout, we use $k$ to denote the number of ancestors (i.e. the number of clades) at the cut-off time. |
| $X_{k'}$ | The set of possible partitions after $k' = n - k$ coalescent events. Throughout, we use $k'$ to denote the number of coalescent events by the cut-off time. |
| $\mathcal{A}(x) = (\alpha_1, \dots, \alpha_n)$ | The “clade representation” of the partition $x$. Here $\alpha_j$ gives the number of clades in the partition $x$ of size $j$ ($1 \leq j \leq n$). |
| $\mathcal{B}(C) = (\beta_1, \dots, \beta_n)$ | The “clade representation” of the tuple $C$. Here $\beta_j$ is the number of clades in the tuple $C$ of size $j$ ($1 \leq j \leq n$). |
| $l$ | A subscript indicating the number of “sub-sampled” lineages. |
| $y_l^C = (i_1, \dots, i_c)$ | A “lineage configuration” is a tuple representing the number of the $l$ “sub-sampled” lineages belonging to each of $C$ clades of size $C = (m_1, \dots, m_c)$. Here $1 \leq i_j \leq m_j$ is the number of lineages belonging to the clade of size $m_j$ ($1 \leq j \leq c$). |
| $Y_l^C$ | The set of all lineage configurations ($y_l^C \in Y_l^C$) of $l$ randomly “sub-sampled” lineages (sampled without replacement) from a set of clades $C$. |

Below we calculate the closed-form expressions for five probability mass functions. Each represents either the probability of sampling a clade, or set of clades, of a specific size(s) (throughout we refer to the “sampling” of clades) or the probability of “sub-sampling” lineage(s) within a clade(s) of specific size(s) (throughout we refer to the “sub-sampling” of lineages) given a cut-off time *T* and a total of *n* observed lineages (see Table 2). In Case 1, we consider the probability that a randomly sampled clade has size *m* (*m* ≤ *n*). Case 2 considers an alternative sampling scheme, considering the probability that a randomly sub-sampled lineage belongs to a clade of size *m* (*m* ≤ *n*). Case 3 complements Case 1 by considering the joint probability that *c* randomly sampled clades have sizes (*m*_1_, …, *m*_*c*_) where clades are sampled without replacement (of the clade). In Case 4, we consider the probability that *l* randomly sub-sampled lineages drawn without replacement belong to *c* distinct clades of size (*m*_1_, …, *m*_*c*_). Finally, Case 5 is a generalization of Case 1 that considers the probability that a randomly sampled clade has size *m* given that the number of niches (aka the “niche size”) varies through time as given by *N*(*τ*) at time *τ* in the past measured in coalescent time units of *N*_ref_ generations. Although not presented here, a similar generalization of Cases 2-4 for non-constant niche sizes can be derived.

**Table 2:**
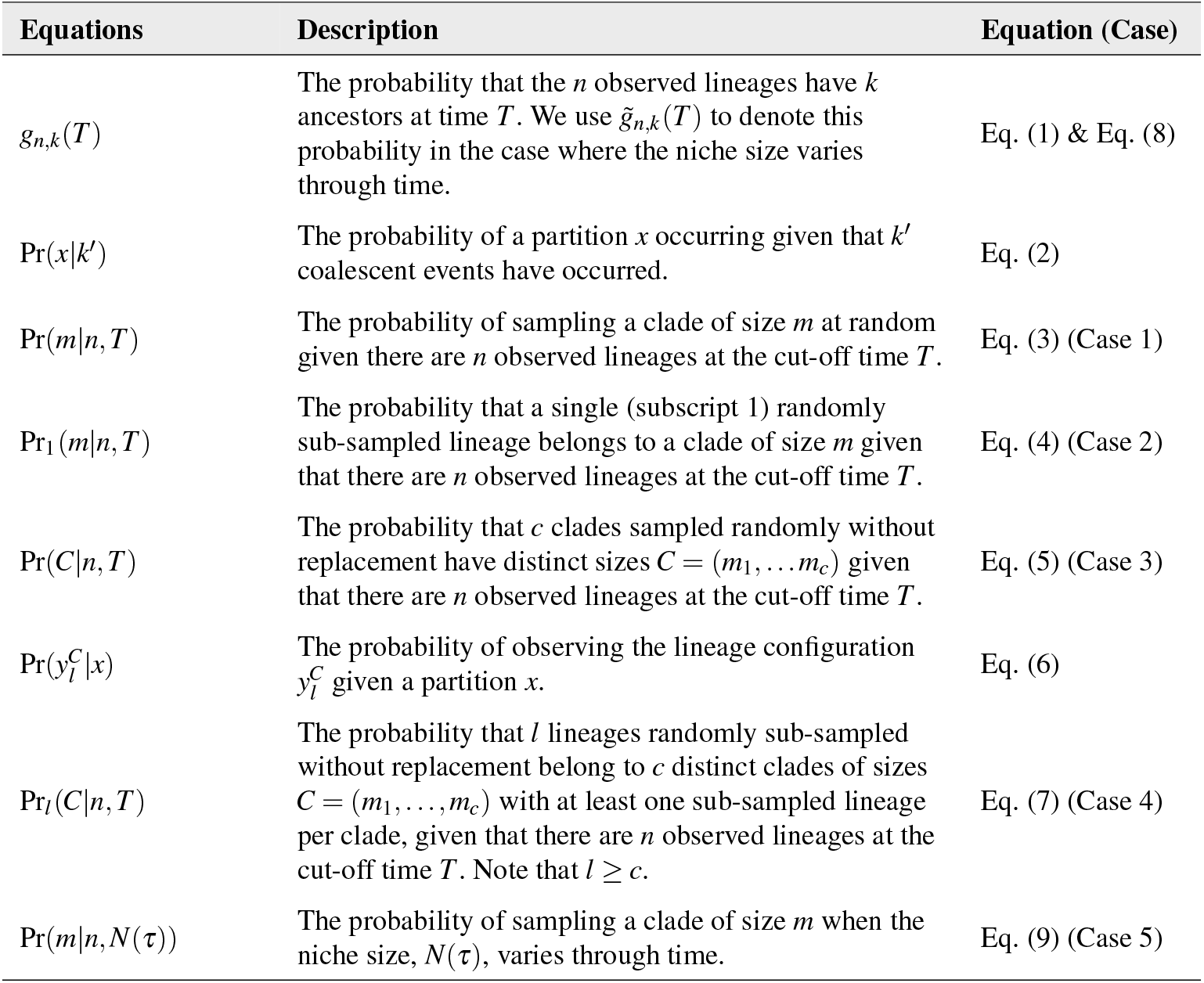
List of probability distributions.

Regardless of the case considered, we derive the desired probabilities using a three-step process outlined in Fig. 2. We begin (Step 1 in Fig. 2) by considering the probability that there are *k* ancestors at the cut-off time *T*, or in other words, the probability that *k*′ = *n* − *k* coalescent events have occurred by time *T*. We denote this probability as *g*_*n,k*_(*T*) (see Table 2). Following Wakeley ([27] Equation 3.41), when the niche size remains constant, *g*_*n,k*_(*T*) is given by:

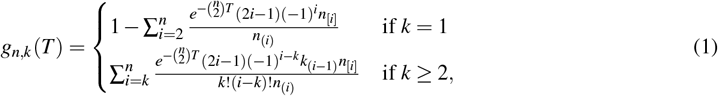

where *n*_[*i*]_ is the descending factorial and *n*_(*i*)_ is the ascending factorial. When the niche size varies through time, this probability must be derived separately (see Case 5 below).

**Figure 2:**
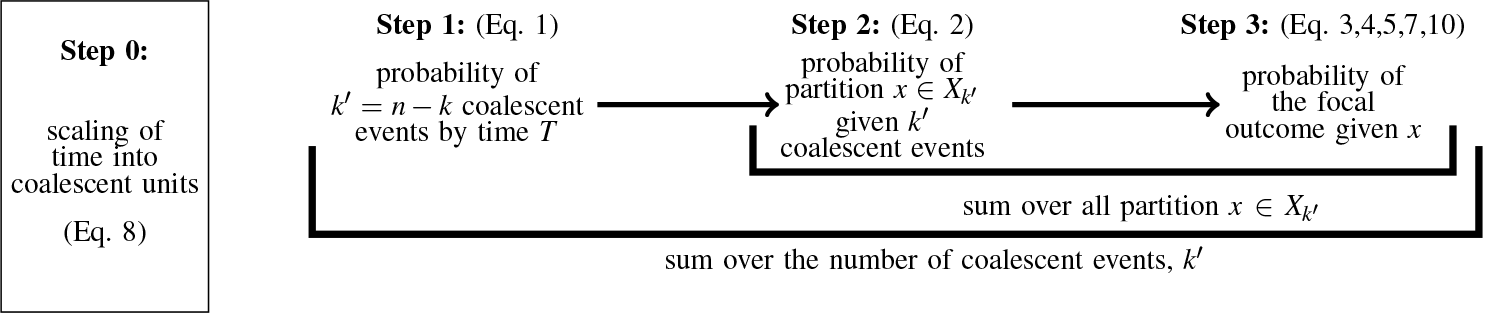
Overview of the derivation of clade size probabilities.

Given that *k*′ coalescent events have occurred by the cut-off time, we then examine the clades formed using the partitions of the integer *n* of size *k* (Step 2 in Fig. 2). It’s worth recalling the notion of an integer partition. A partition *x* of a positive integer *n* of size *k* (hereafter referred to simply as a ‘partition’) is a set of non-decreasing positive integers *x* = *{s*_1_, *s*_2_, …, *s*_*k*_*}* such that 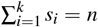.In the coalescent model, the partition *x* represents the clustering of the *n* lineages into *k* clades, each with a distinct ancestor at the cut-off time. For each 1≤ *j* ≤ *n*, let *α*_*j*_ be the number of times the integer *j* appears in the partition *x* such that *α*_*j*_ tells us how many clades are formed of size *j*. We call the list of *α* values for a particular partition *x* the ‘clade representation’ of this partition, which we denote by 𝒜(*x*) = (*α*_1_, *α*_2_, …, *α*_*n*_). 𝒜(*x*) satisfies the constraint ∑ _*j*_ *α*_*j*_ = *k*. Note that each partition, *x*, corresponds to a unique clade representation, 𝒜(*x*), and vice versa.

Using the notion of partitions, given that *k*′ coalescent events have occurred, one or more partitions could arise, with some partitions more likely to occur than others (Fig. 3). The probability of observing a given partition can be calculated using the classic Pólya Urn model to describe the coalescent process [30, 31, 32]. Specifically, balls in the Pólya Urn’s model represent observed lineages (*n* total balls). Urns represent clades such that at the present day, there are *n* urns, each with a single ball (each lineage is in a unique clade). When a coalescent event occurs, we randomly pick two urns, combine the balls into one urn, and remove the now-empty urn. Note that throughout each urn must contain at least one ball. The number of balls *s* in each urn at time *T* forms a partition arising from the *k*′ coalescent events. To describe this process, let *X*_*k*_ be the set of possible partitions that can arise after *k*′ coalescent events given *n* initial lineages. For example, suppose *n* = 6 (we will use the case of *n* = 6 as an illustrative example throughout as shown in Fig. 3). Initially (before the first coalescent event), there is a single possible partition, *X*_0_ = [{1, 1, 1, 1, 1, 1}], with each ball in a separate urn. After a single coalescent event, there is still only one possible partition *X*_1_ = [{1, 1, 1, 1, 2}] with a single pair of balls in a shared urn. After a second coalescent event, however, there are multiple possible partitions, *X*_2_ = [{1, 1, 2, 2}, {1, 1, 1, 3}], note that both of these partitions consist of 4 urns and can be distinguished from one another by their distinct clade representations. The process continues as shown in Fig. 3 until a single partition arises after *n* − 1 coalescent events, *X*_*n*−1_ = [{*n}*].

**Figure 3:**
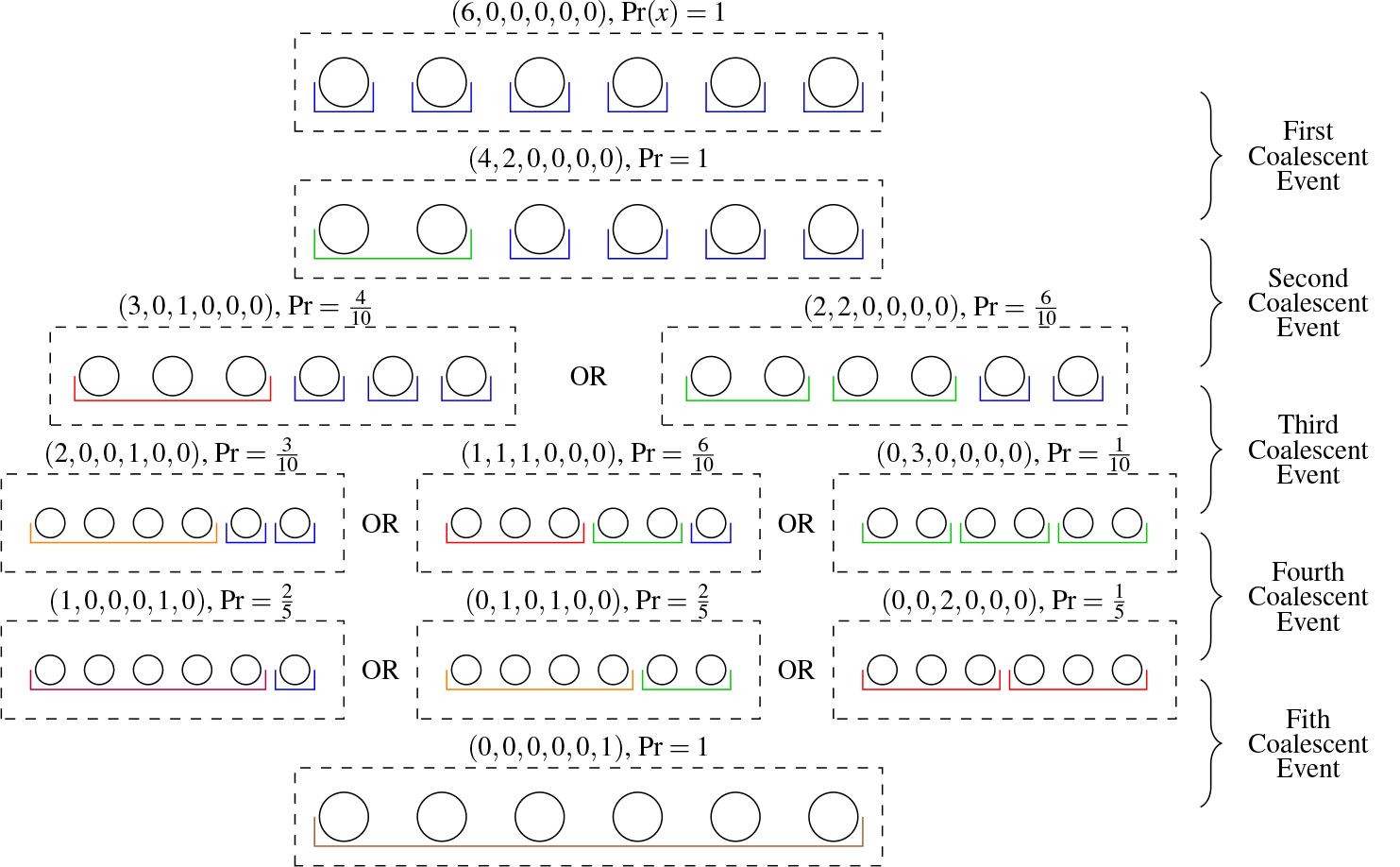
Possible partitions and coalescent events with *n* = 6 samples. Each row lists the possible partitions that could arise following *k*′ = 0, 1 … 5 coalescent events. Circles represent lineages (balls), coloured open boxes represent clades (urns), dashed boxes are partitions, and tuples above each dashed box are that partition’s clade representation, 𝒜(*x*). The probability of the partition can occur given the *k*′ coalescent events are listed beside the clade representation. The number of circles inside the urns represents the clade sizes as denoted by colour (colours are used throughout).

If there is more than one partition after *k*′ coalescent events, the probability that each partition arises from the Wright-Fisher process may differ. For example, the probability of {1, 1, 2, 2} and {1, 1, 1, 3} occurring after two coalescent events are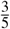 and 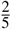, respectively. Hence, in Step 2, we calculate the probability of obtaining a particular partition *x* ∈ *X*_*k*′_.

### Theorem

The probability of obtaining a partition *x* ∈ *X*_*k*′_ given *k*′ coalescent events have occurred (i.e., that there are *k* total clades) with clade representation 𝒜(*x*) through the Wright-Fisher process is given by:

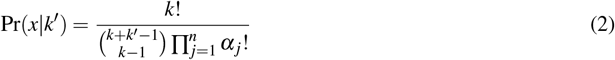

where ∑ _*j*_ *α*_*j*_ = *k*.

*Proof:*

In summary, we calculate this probability by considering the number of ways of rearranging *k* urns divided by the total number of possible configurations of *n* balls among the *k* urns, given the constraint that each urn must contain at least one ball. The number of ways that all the urns of the same size are indistinguishable is:

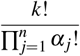

It remains to count the total number of possible configurations of *n* balls among the *k* urns, given the constraint that each urn must contain at least one ball. Namely, this constraint requires *k* balls to be placed one in each urn with *n* − *k* = *k*′ remaining balls distributed among the urns. The total number of ways of distributing these *k*′ balls among the *k* urns is:

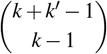

Hence, the probability of a given clade representation is as noted in the theorem. □

To complete the derivation, we consider the probability of obtaining the desired sampled clades or sub-sampled lineages from a particular partition (Step 3 in Fig. 2) for each Case described below.

*Case 1:*

Here, we consider the probability that a randomly sampled clade has size *m* given the number of observed lineages, *n*, and cut-off time, *T*. To complete the derivation (Step 3 in Fig. 2), we consider the proportion of clades formed of the target size, *m*, for each partition, *x*. In this case, this is 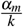. Summing over all the number of possible coalescent events *k*′ and the possible partitions given that *k*′ coalescent events have occurred, *x* ∈ *X*_*k*′_, and weighting accordingly we obtain:

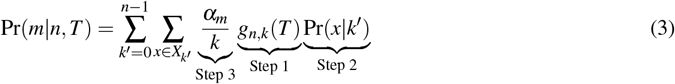

where *k* = *n*−*k*′. Substituting Eq. (1) and (2) into Eq. (3) and simplifying, we obtain:

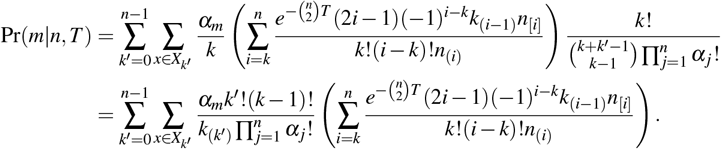

*Example n* = 6 & *m* = 3:

There are in total 11 possible partitions that could occur given *k*′ ={ 0, 1, …, 5} coalescent events (see boxes in Fig. 2), of which only three contain a clade of the focal size *m* = 3: *x*_2_ = {1, 1, 1, 3 }, *x*_3_ = {1, 2, 3 }, *x*_4_ = 3, 3 where the subscript here denotes the number of coalescent events that have occurred. Using Eq. 3, the probability of sampling a clade of size 3 given 6 observed lineages and a cut-off time *T* is:

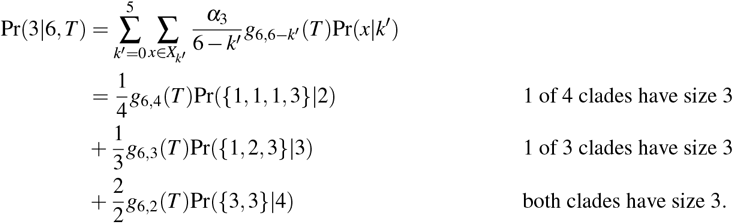

*Case 2:*

Here, we consider the probability that a single sub-sampled lineage belongs to a clade of size *m* given the number of observed lineages and cut-off time. Following Step 3 (Fig. 2), for the partition *x*, the probability that a randomly sampled lineage belongs to a clade of size *m* is: 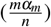.We obtain the desired probability by summing over the possible partitions and coalescent events, as above.

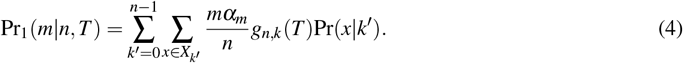

An example of *n* = 6 & *m* = 3 is given in Appendix D.1.

*Case 3:*

Here, we consider the probability that *c* (1≤ *c*≤ *k*) clades, sampled randomly without replacement, have sizes *C* = (*m*_1_, *m*_2_, …, *m*_*c*_) given the total number of observed lineages and cut-off time. We denote the desired clade sizes with the tuple *C* = (*m*_1_, …, *m*_*c*_) where two or more elements of the tuple (e.g., *m*_*i*_ and *m*_*j*_) may be the same. To facilitate the calculation, we define the ‘clade representation’ of the tuple ℬ(*C*) = (*β*_1_, …, *β*_*n*_) where *β*_*j*_ is the number of clades in the tuple *C* with size *j* (see Figure 1). The probability of randomly sampling the *c* clades of the desired sizes equals:

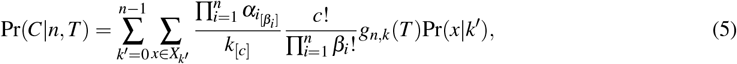

where 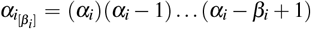 and *k*_[*c*]_ = (*k*)(*k* − 1) … (*k* − *c* + 1) are the descending factorials. An example of *n* = 6 & *m*_1_ = *m*_2_ = 2 is given in Appendix D.2.

*Case 4:*

Here we consider the probability that *l* random lineages sub-sampled without replacement belong to *c* clades of sizes *C* = (*m*_1_, …, *m*_*c*_) (where *l*≤ *c*) given the total number of observed lineages, *n*, and cut-off time, *T*. As with Case 3, we denote the clade representation of the tuple *C* with ℬ(*C*) = (*β*_1_, …, *β*_*c*_). To facilitate the calculation, we define a ‘lineage configuration’ of the *l* lineages among the sampled clades 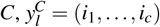, where *i* _*j*_ is the number of the lineages sub-sampled from the *j*^*th*^ clade which has size *m*_*j*_. We denote the set of possible lineage configurations as 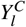. Note that these lineage configurations are subject to the constraint that *l* _*j*_≤ *m*_*j*_.

To complete Step 3 (Fig. 2), we calculate the probability of observing the lineage configuration 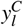 given a partition *x*. We have:

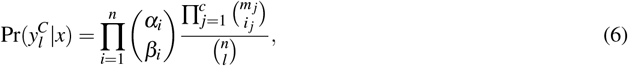

where *α*_*j*_ denotes an element of the clade representation of the partition *x* and *β*_*j*_ an element of the clade representation of the sampled clades *C*. Here 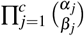 is the number of ways of choosing the desired clades and 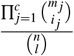 is the probability of observing the lineage configuration from lineages among the desired clades. Completing the derivation as above, we have:

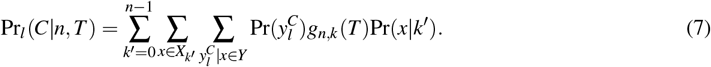

An example of *n* = 6 & *m*_1_ = 1, *m*_2_ = *m*_3_ = 2 & *l* = 4 is given in Appendix D.3.

*Case 5:*

Here, we consider the probability of sampling a random clade of size *m*, allowing the niche size, *N*(*τ*), to vary through time. To do so, we first calculate the probability that there are *k* ancestors of the *n* observed lineages at time 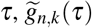 (See Appendix B and Step 1 in Fig.2):

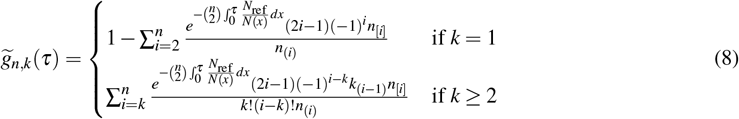

Given the probability 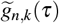, the probabilities in Steps 2 and 3 (Fig. 2) are independent of niche size, and hence the probability of sampling a clade of size *m*, Pr(*m*|*n, N*(*τ*)) equals:

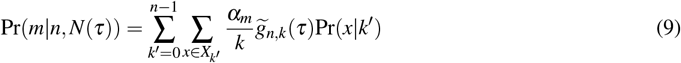

While the result in Eq. 9 can apply for any niche size trajectory, for numerical purposes, we consider the specific case where the niche size grows exponentially at rate *r* according to *N*(*τ*) = *N*_0_*e*^−*rτ*^ (i.e., a declining niche size backward in time). In this case, there exist three logical choices for the reference niche size *N*_ref_ that would allow for straightforward comparisons of the distribution of clade sizes between an exponentially growing versus constant niche size model to calculate the scaled time 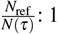) the reference size equals the size at the present day: *N*_ref_ = *N*_0_, 2) the reference size equals the arithmetic-mean niche size between the present day and the cut-off time: 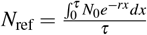, and 3) the reference size equals the harmonic-mean niche size between the present day and the cut-off time: 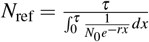. We derived the expressions for the scaled time in Appendix C.

### Validation

We validate our analytical expressions using simulations for each case described above. To do so, we simulate 50, 000 coalescent trees with *n* = 6 observed lineages. We then analyze the resulting (sub-)sampled (lineages) clades at 500 equally spaced cut-off times between *T* = 0 (the present day) and *T* = 3. The validation of Case 1 is shown in Fig. 4. Results for Cases 2-5 are shown in supplementary Figs. E.7-E.10. Simulations were implemented in Python 3.10 and can be found on [GitHub](https://github.com/Yexuan-Song/Clustering/tree/main).

**Figure 4:**
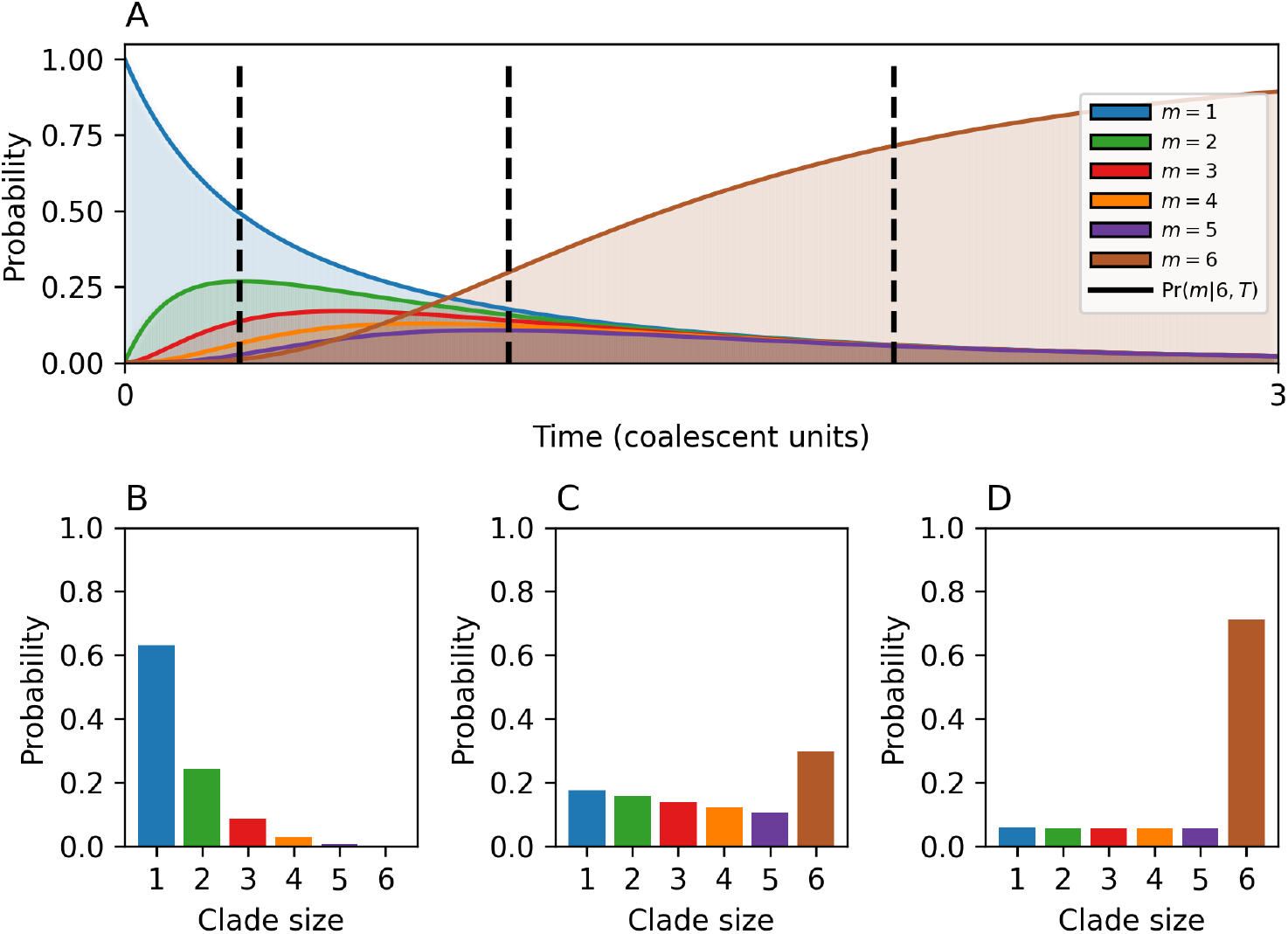
Validating Eq. 3 with simulated data. Comparison of analytical (Eq. 3) and simulated results for the probability of observing a clade of size *m* = {1, 2, … 6} (as indicated by colour) for the case where *n* = 6. Top panel: Clade size over time. Solid lines give analytical predictions. Shaded areas give results for simulations. Bottom panels: Clade size distribution at three exemplary time points (black dashed vertical lines from top panel), *T* = 0.3, 1, &2.

## 3. Results

Here we analyze the (joint) distribution of clade size(s) and their biological implications by comparing these distributions to either the expectations in the birth-death diversification model (e.g., Case 1) or by comparing the distributions in each case to one another. We begin by comparing the probability of observing a random clade of size *m* in the coalescent model (Case 1) to expectations in the density-independent birth-death [3] or Yule (birth only) [1] model. Known as the “hollow curve”, clade size in the birth-death family of diversification models is expected to follow a geometric distribution with the probability decreasing as clade size *m* increases. As the maximum clade size is unconstrained in the birth-death model but limited to a maximum size *n* in the coalescent model, a quantitative comparison is not straightforward. Nevertheless, we can compare the results qualitatively to the “hollow curve” by considering the distribution of clade sizes at three characteristic time points (Fig. 4B-D). We find that, near the present day (Fig. 4B), the probability of observing a clade of size *m* decreases monotonically with increasing *m* in an approximately geometric manner analogous to the “hollow curve”. Deeper in the tree (Fig. 4C), however, subsequent coalescent events increase the probability of observing larger clades, with clade size now declining nearly linearly between *m* = 1 and *m* = *n* − 1 with an overabundance of clades of size *n* arising as the cut-off time exceeds the T_MRCA_ of the *n* observed lineages. This process continues until the cut-off exceeds the T_MRCA_ in the vast majority of cases (Fig. 4D), with the remaining clade sizes occurring with equal probabilities. While we have focused on *n* = 6 throughout, a similar pattern is observed for larger *n* (see Fig. E.11 for *n* = 30). One key advantage of examining coalescent processes is the existence of analytical expressions for both the marginal and joint clade size distributions, sampling of two or more clades, or sub-sampling of two or more lineages from within clades. In the coalescent model, there are two sources of non-independence when (sub-) sampling (lineages) clades. First, there is the inherent density-dependence in the diversification process due to the fixed niche size. Second, there is the constraint of having a fixed number of observed lineages “used up” when more than one lineage is sampled. The relative contributions of these two forms of non-independence can be intuited through comparisons of the five cases above.

First, consider the comparison between the random sampling of a single clade of size *m* (Case 1) and the random sub-sampling of a lineage from within a clade of size *m* (Case 2) (Fig. 5A). Sub-sampling lineages down-weights the probability of observing a singleton clade (*m* = 1) and up-weights the probability of observing a clade of intermediate size (1 *< m < n*). Finally, the probability of obtaining a clade of size *m* = *n* (i.e. all observed lineages belong to a single clade) is independent of whether one samples a clade or sub-samples a lineage within a clade, as both cases only depend on whether the cut-off time is greater than or equal to the time to the most recent common ancestor *T*≥ *T*_*MRCA*_. The reduced probability of sampling a clade of intermediate size (and the equal and opposite increase in the probability of sampling a clade of size *m* = 1) captures the non-independence due to fixed *n* as multiple lineages are “used up” when sampling clades, but only a single lineage is used when sub-sampling lineages within these clades.

**Figure 5:**
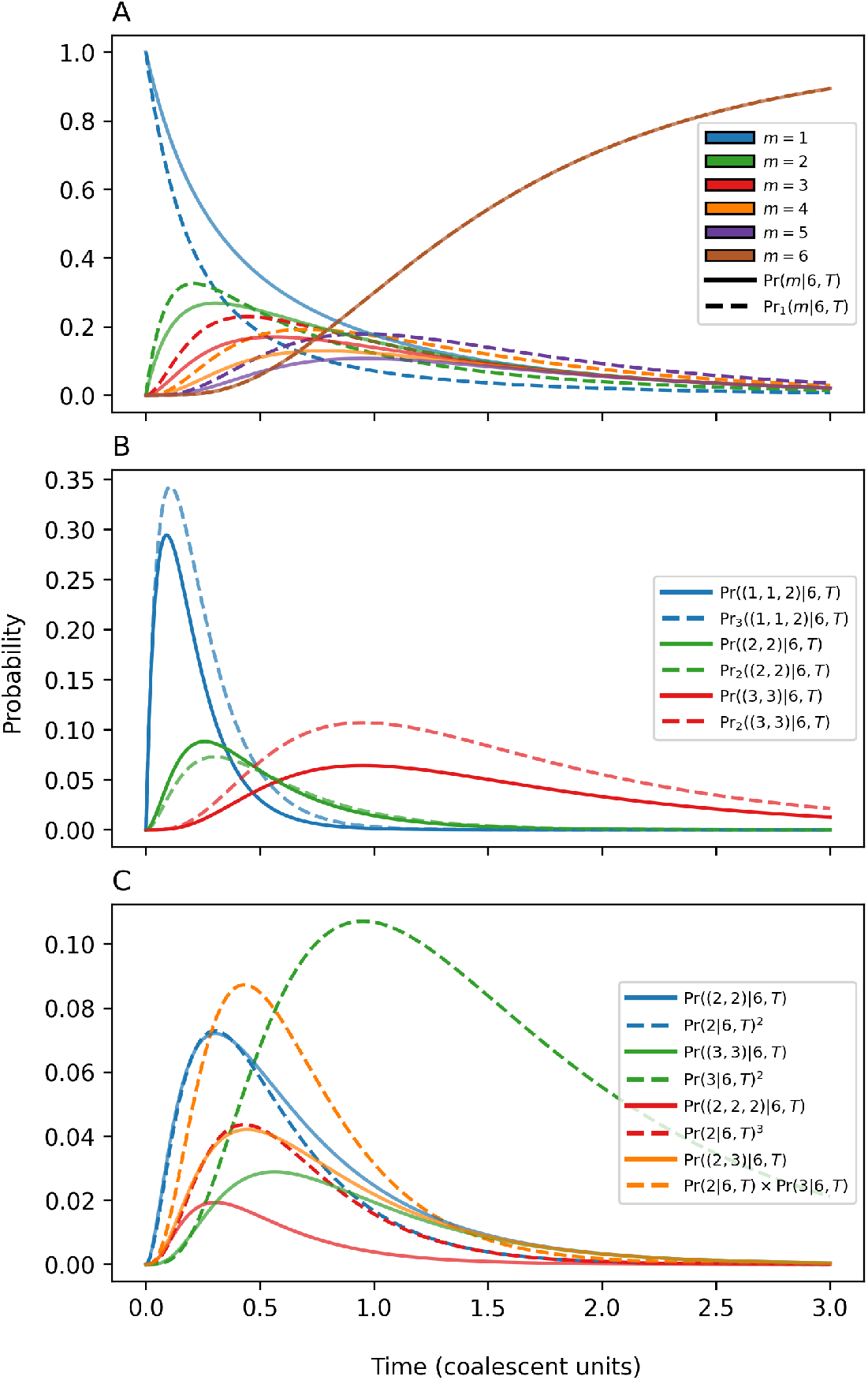
Marginal and joint clade size distributions. Comparison of the distribution of clade sizes (heights of the coloured curves at a given time) as a function of the cut-off time *T* for cases with *n* = 6 observed lineages. A: Comparison of Cases 1 and 2: the probability of sampling a single clade of size *m* by either sampling a clade (solid) or sub-sampling a lineage from the clade of size *m* (dashed). B: Comparison of Cases 3 and 4 of sampling multiple clades (solid) at random or sub-sampling lineages (solid). C: Comparison of the joint probability multiple clades (Case 3, solid) relative to expectations from the marginal distribution assuming independence (Case 1, solid).

The net effect of a fixed number of observed lineages is more complicated when comparing the sampling of multiple clades (Case 3) and sub-sampling of multiple lineages from within clades (Case 4) (Fig. 5B). In this case, removing lineages/clades with sampling can either decrease the probability of obtaining the desired clade sizes when sub-sampling lineages (red and blue cases) or increase it (green case). To understand this, consider the red curves in Fig. 5B, where one wishes to (sub-)sample two clades of size 3. Here, we are interested in only a single partition *x* = {3, 3}. When sub-sampling lineages, Pr_2_((3, 3) | 6, *T*), we are considering the probability of sub-sampling two lineages, one from each clade of size 3. In comparison, Pr((3, 3) | 6, *T*) is the probability of sampling both clades with removal. The probability of (sub-)sampling the first (lineage) clade of size 3 is independent of the sampling design. However, in the sub-sampling case, only a single lineage is removed such that 3 of the 5 remaining lineages belong to the second clade of size 3. In comparison, when the first clade is removed following sampling, we are guaranteed to sub-sample the second clade of size 3. Hence, the net effect of the sampling scheme (Case1-4) depends on (sub-) sampled clades and the need to obtain distinct clades (see Fig. 5B-C).

Finally, the effect of non-independence due to a fixed niche size/density-dependence can be intuited by comparing a model in which the niche size remains constant to one in which it grows exponentially. Figure 6 shows one such comparison where the constant/reference niche size equals the arithmetic mean niche size. In this case, the arithmetic mean niche size is larger than the “effective niche size,” which, from population genetics theory, is known to be the harmonic mean. Consequently, coalescent events occur faster in the variable niche size model, favouring intermediate clade sizes in the short term and clades of size *m* = *n* in the long term.

**Figure 6:**
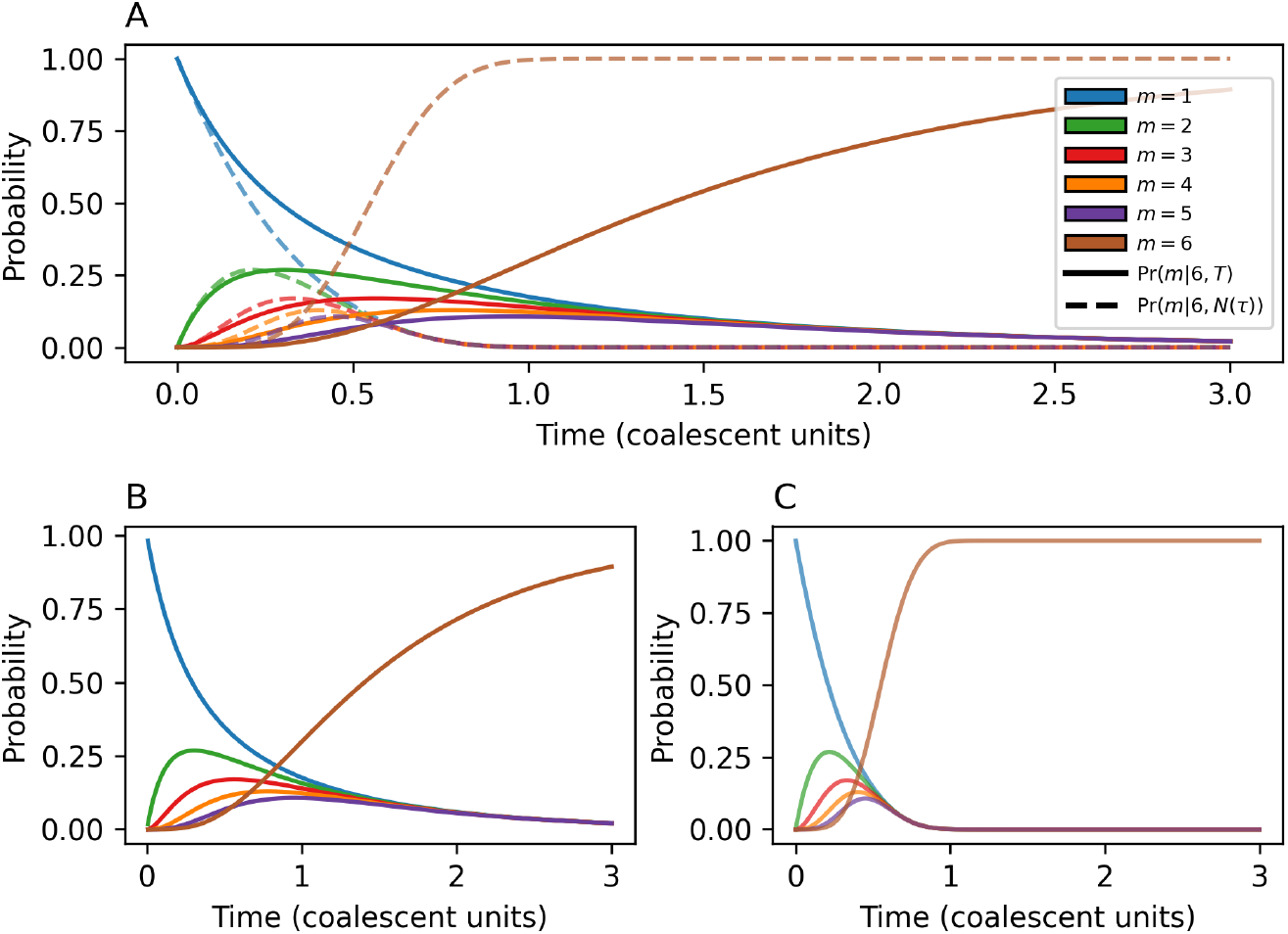
Comparison of clade size in an exponentially growing versus a constant niche size model. The distribution of clade sizes over time given a constant niche size *N*_ref_ (solid) versus in an exponentially growing niche size model as given by Eq. 3 and 9, respectively. A: Reference size of the arithmetic mean. B: Reference size of the harmonic mean. C; Reference size is the niche size at the present day.

## 4. Discussion

Here we have derived analytical expressions for the marginal and joint clade size distributions under the coalescent diversification model. These distributions complement existing measures of diversity (e.g., Ewens’s Sampling Formula and the Site Frequency Spectrum) that are grounded in applications of the coalescent to population genetics. Methodologically speaking, a key insight of this work is that clades in the coalescent model can be represented as integer partitions of the total number of observed lineages, *n*. As in other applications of the coalescent as a tree prior, the resulting analytical tractability allows us to derive exact expressions for a model in which diversification is strongly density-dependent. Hence these results provide a useful alternative to expectations for clade size in the birth-death(-sampling) model [1, 3]. By qualitatively comparing the results from the coalescent diversification model to that of the birth-death model, we find that the two models produce similar expectations (the “hollow-curve”) for only young clades before diversity patterns in the coalescent are limited by the number of observed lineages.

One key attribute of the coalescent process is that it can be used to study diversification when the niche size dynamics, *N*(*τ*), vary in a pre-determined way. As such, the clade size distributions developed here (particularly in Case 5) can be used as the basis for testing between alternative diversification hypotheses; for example, a diversification model in which the number of niches remains constant through time versus one in which niche size grows exponentially [14, 15]. Similarly, the probability distributions developed here can also be used to identify diversity outliers, speciose or depauperate clades [8, 9, 10, 11]. In particular, the joint-clade size distributions may be useful in performing sister-clade comparisons.

Throughout we have referred primarily to diversification within a macroevolutionary context, e.g., referring to ‘niches’ and ‘species’. However, the diversification of viruses or other pathogen populations can result in similar evolutionary histories. Within this epidemiological context, the niche size, *N*(*τ*), represents the number of infected hosts and the dynamics of the epidemic through time. Similarly, clades in this context represent transmission ‘clusters’, and ‘species’ are sampled pathogen sequences. Used widely in outbreaks of Human immunodeficiency virus (HIV) infection [33, 34, 35, 36, 37, 38, 39], the clustering of viral samples can be used to identify high-risk groups and to study the role of co-variates (e.g., co-infection with Hepatitis C virus [40]) on transmission. The coalescent diversification model may be particularly well suited to epidemiological applications where the number of sequenced samples is limited relative to the total outbreak size (e.g., *n*≪ *N*). Similarly, the ability to study clade size in a growing, constant, or declining epidemic (Case 5) allows us to link the size of observed clusters to the underlying epidemiological dynamics.

While the analytical expressions derived above theoretically apply for all *n* ≪ *N*, practically speaking, it is only computationally feasible to evaluate these expressions for trees with on the order of 100 tips. The computational feasibility problem arises from the property of integer partition. The number of partitions, *p*(*n*), grows exponentially, with a rate proportional to the square root of the sample size *n* [31]:

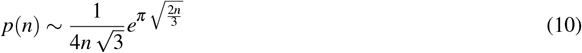

For example, if there are *n* = 1000 lineages, the total number of partitions is approximately *p*(1000) ≈ 2.4*×*10^31^. Application to large trees (e.g., a pandemic with substantial sampling effort) would necessitate finding an approximation to the analytical solutions when *n* is large. There also exist conceptual limitations when applying clade size to the analysis of epidemiological clusters. Despite the substantial attention devoted to identifying transmission clusters, there is no universal agreement on how clusters should be defined [33, 34]. Clusters are sometimes based on genetic distances [35, 37] while others are clustered based on phylogenetic distances [38, 39]. Bridging these definitions would require incorporating mutation into the coalescent process described above. Nevertheless, the expectations derived here could provide the foundation for testing when and how the accuracy of cluster identification changes with mutation rates and epidemiological dynamics.

In conclusion, our study provides an analytical framework for understanding clade size distributions under the coalescent model. We propose various sampling schemes and extend the model to account for niche dynamics. We propose future directions for hypothesis testing and model fitting in macro-evolution and epidemiological studies.

## Acknowledgments

This work is supported by NSERC (AM: CRC-2021-00276 and RGPIN-2022-03113, CC: Canada’s 150 Research Chair program and RGPIN-2019-06624).

## Appendix A. Availability of materials

Codes can be accessed at the following GitHub repository.

## Appendix B. Derivation of 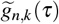

Here we consider the probability of observing *k*′ = *n* − *k* coalescent events by time *τ* under the coalescent model, where the niche size *N*(*τ*) is allowed to vary through time. Time *τ* is measured backward-in-time in coalescent units of *N*_ref_ generations (see Table 1). Given the reference niche size *N*_ref_ we define a scaled niche size 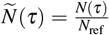, see Griffiths and Tavaré [41] and Appendix C. The derivation below follows the same structure as [42].

Given this scaled niche size, we consider a continuous-time discrete-state Markov process *A*(*τ*) following the number of sampled ancestors present at time *τ* before the present day, assuming there are *n* sampled lineages at the present day: *A*(*τ*) = *{A*_*τ*_ = *k*|*A*_0_ = *n, Ñ* (*τ*)*}*. As sampled lineages can only coalesce, reducing the number of ancestors, *A*(*τ*) is a “death process” which has a time-dependent transition rate matrix 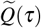 which is a lower bidiagonal matrix of the form:

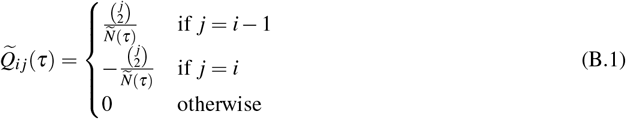

The common denominator *Ñ*(*τ*) can be factored out and gives 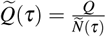, where the resulting matrix *Q* is time independent.

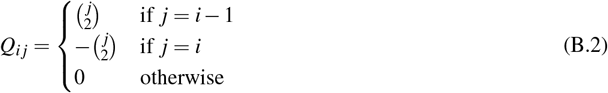

Using the Chapman-Kolmogorov equation, we can derive the time-dependent transition probability matrix 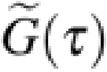. Where the *n, k*-th element of this matrix 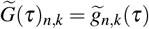 is the desired probability of transitions from *n* sampled ancestors at the present day (*A*_0_ = *n*) to *k* sampled ancestors at time *τ* in the past *A*_*τ*_ = *k*.

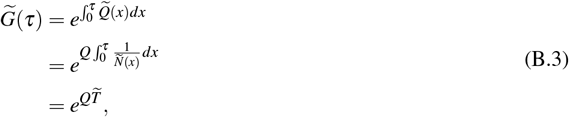

where 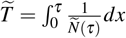 is the scaled time.

To calculate the matrix exponential, we first find the eigendecomposition of 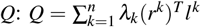. Where *λ*_*k*_ is the *k*th eigenvalue and *r*^*k*^ and *l*^*k*^ are, respectively, the right and left eigenvectors corresponding to the *k*th eigenvalue *λ*_*k*_. To find the eigendecomposition of the matrix *Q*, it is helpful to note the form of the matrix *Q*:

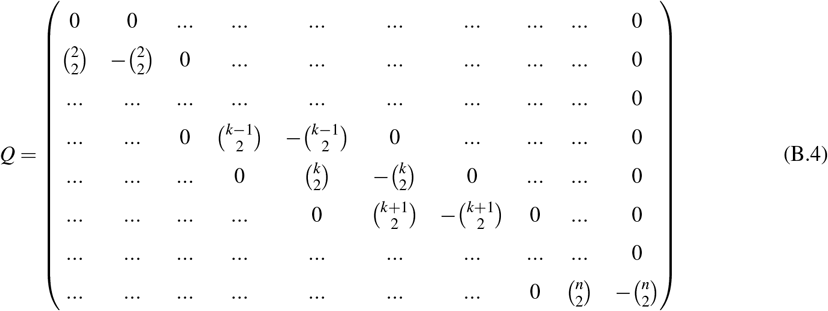

First, given its form, the eigenvalues are given by the diagonal values: 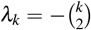. Given these eigenvalues, we find the corresponding left and right eigenvectors element by element. Let 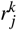 and 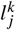 be the *j*th element of the right and left eigenvector corresponding to the eigenvalue *λ*_*k*_, respectively. To find *r*^*k*^, we solve the equation (*Q* − *λ*_*k*_*I*)*r*^*k*^ = 0. The solution gives the following recurrence relation: 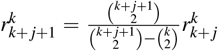 and 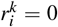 for *i < k*. Similarly, to find *l*^*k*^, we solve the equation (*l*^*k*^)^*T*^ (*Q*−*λ*_*k*_*I*) = 0. The solution gives the following recurrence relation: 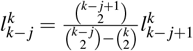 and 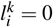 for *I* > *k*. Using induction, we can show:

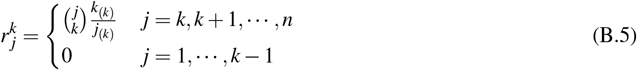

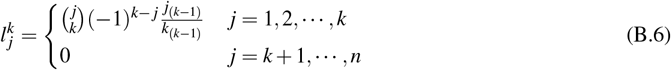

We can calculate the desired probability by considering the *n, k* element of the transition probability matrix: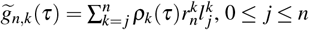. where 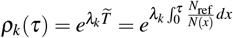. This reduces to the result in Eq. 8.

## Appendix C. The derivation of scaled time, 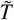

Here, we use different reference niche size *N*_ref_ choices to find the scaled time 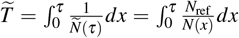 described in Eq. 8, where *N(x)= N 0e*^−*rx*^. We propose the following three choices: (1). *N* _ref_ = *N*_0_, (2). 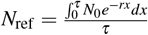, and (3). 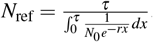.

1. Reference size to the present day (*N*_ref_ = *N*_0_):

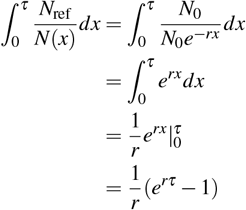
2. The arithmetic mean size between the present day and the cut-off time 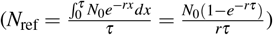:

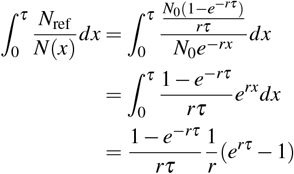
3. The harmonic mean size between the present day and the cut-off time 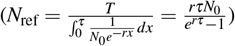:

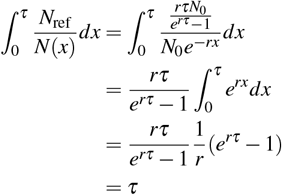

The harmonic mean reference size rescales the time *τ* back to the constant case.

## Appendix D. Examples for different sampling schemes (Case 2 - 4)

### Appendix D.1. Case 2

*Example n* = 6 & *m* = 3:

Using Eq. 4, we can calculate the probability that a randomly sub-sampled lineage belongs to a clade of size 3 given there are 6 observed lineages and a cut-off time *T*.

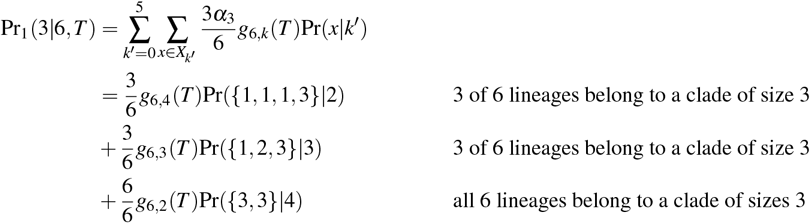

### Appendix D.2. Case 3

*Example n* = 6 & *m*_1_ = *m*_2_ = 2:

Using Eq. 5, we can calculate the probability of observing 2 randomly chosen clades have both sizes *C* = (2, 2).

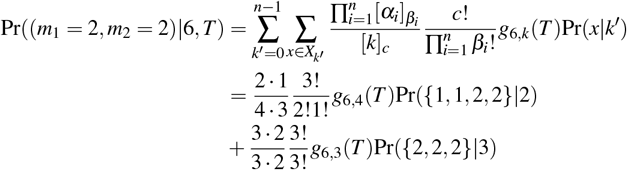

### Appendix D.3. Case 4

*Example n* = 6 & *m*_1_ = 1, *m*_2_ = *m*_3_ = 2 & *l* = 4:

Using Eq. 7, we can calculate the probability of observing 4 sub-sampled lineages, *l* = 4, sampled without replacement, belonging to *c* = 3 clades of sizes *C* = (1, 2, 2) given that there are a total of *n* = 6 observed lineages.

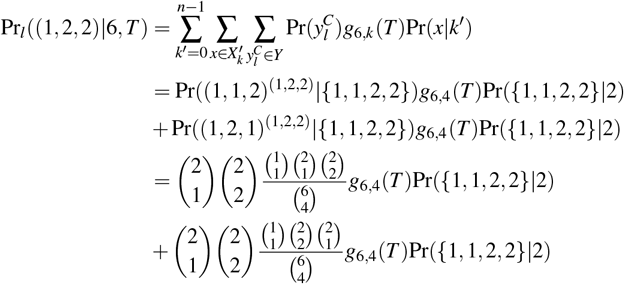

**Table D.3:** Classic diversification models and their attributes. Here, we list whether diversification models are “generative” (they specify an algorithm for simulating a phylogenetic tree) and/or are “evolutionary” (the events in the generative algorithm have evolutionary interpretations [44]) as well as what is (un)known about the three tree attributes: size, shape and branch lengths. The coalescent model is “pseudo-evolutionary” in that it describes the genealogical process and can be used as an approximation to species diversification under strong density dependence.

| Model | Algorithm | Clade Size | Branch Length | Tree Shape |
| --- | --- | --- | --- | --- |
| Yule [1] | Generative / Evolutionary | Known [1] | Known [20] | No |
| Birth-Death(-Sampling) [43] | Generative / Evolutionary | Known [2, 3] | Known | No |
| Coalescent [22] | Generative / Pseudo-Evolutionary | This Study | Known [22] | No |
| β-Splitting [16] | Non-Generative / Not Evolutionary | NA | NA | Yes |
| 2 parameter β-Splitting [44] | Generative / Evolutionary | Unknown | Unknown | Yes |
| PDA [45] | Non-Generative / Not Evolutionary | NA | NA | Yes |
| Multitype Birth-Death [18, 20] | Generative / Evolutionary | Unknown | Known[20] | Yes |

**Table D.4:**
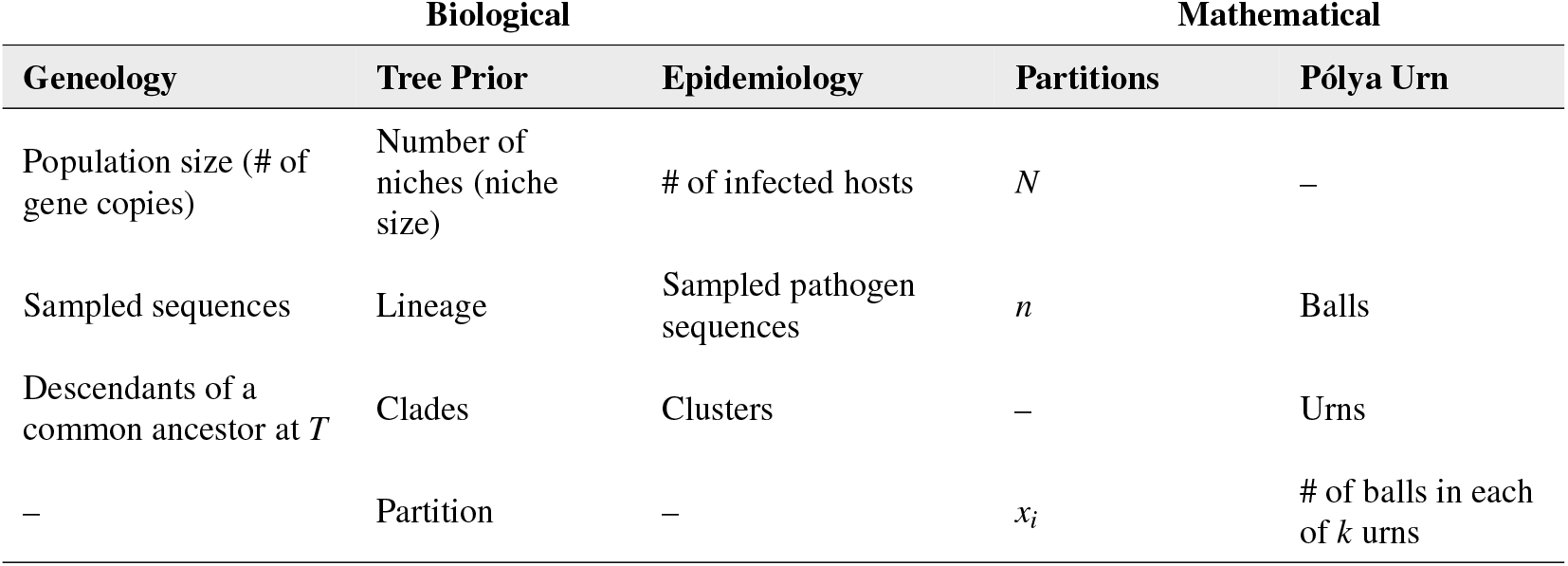
Various terminologies for the components of the coalescent process. Biologically speaking, the coalescent can be used as either a model of an observed gemology or as a tree prior. Mathematically, in this manuscript, we variously refer to the components of the coalescent in terms of integer partitions and in reference to the Pólya Urn model.

| Biological |  |  | Mathematical |  |
| --- | --- | --- | --- | --- |
| Geneology | Tree Prior | Epidemiology | Partitions | Pólya Urn |
| Population size (# of gene copies) | Number of niches (niche size) | # of infected hosts | $N$ | – |
| Sampled sequences | Lineage | Sampled pathogen sequences | $n$ | Balls |
| Descendants of a common ancestor at $T$ | Clades | Clusters | – | Urns |
| – | Partition | – | $x_i$ | # of balls in each of $k$ urns |

## Appendix E. Figures

**Figure E.7:**
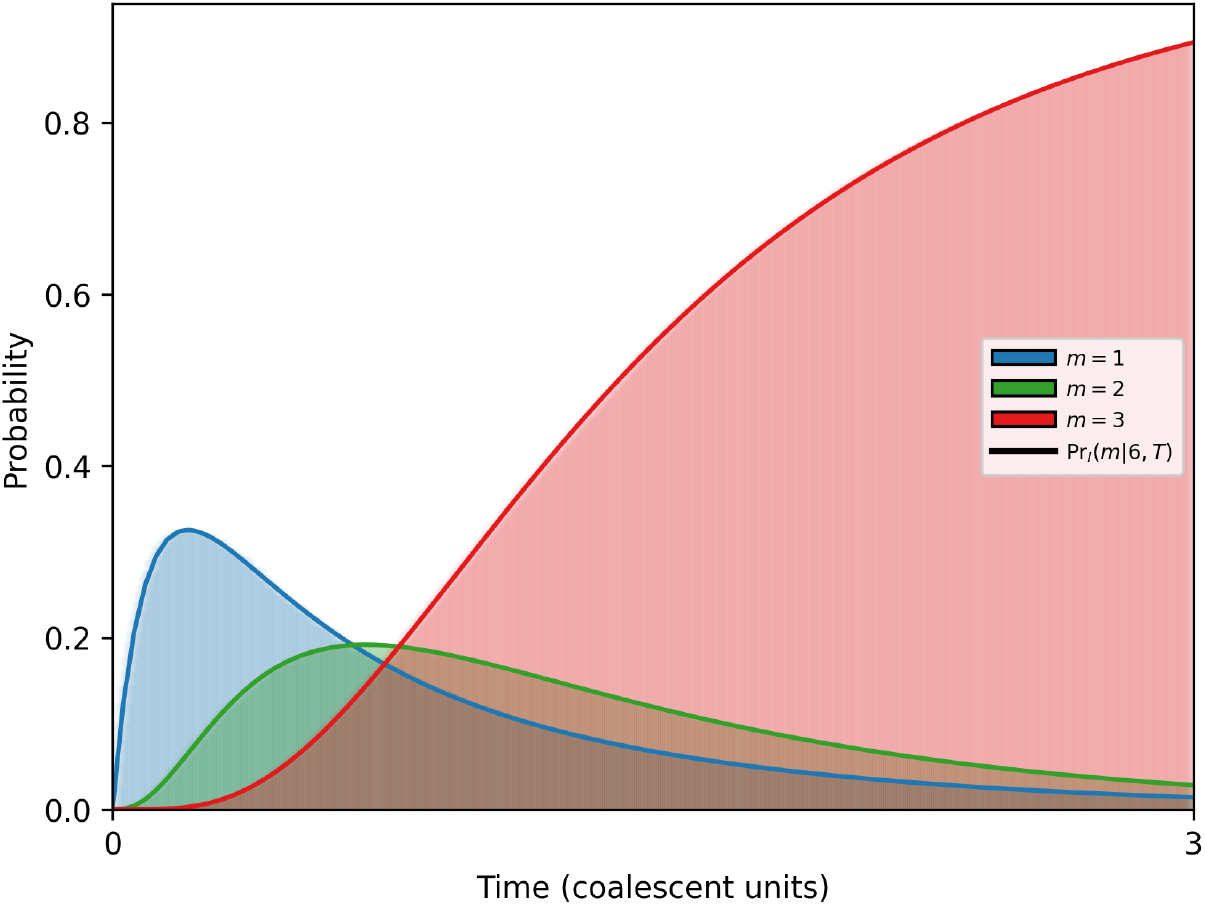
Validating Eq. 4 with simulated data. Here, we simulate the coalescent process and corresponding clade distribution for an example of *n* = 6 sampled lineages. For each time of 500-time points between *τ* = 0 and *τ* = 3, we calculate the probability that a randomly chosen individual belongs to a clade size *m* as shown in the histogram. Solid curves give analytical expectation from Eq. 4.

**Figure E.8:**
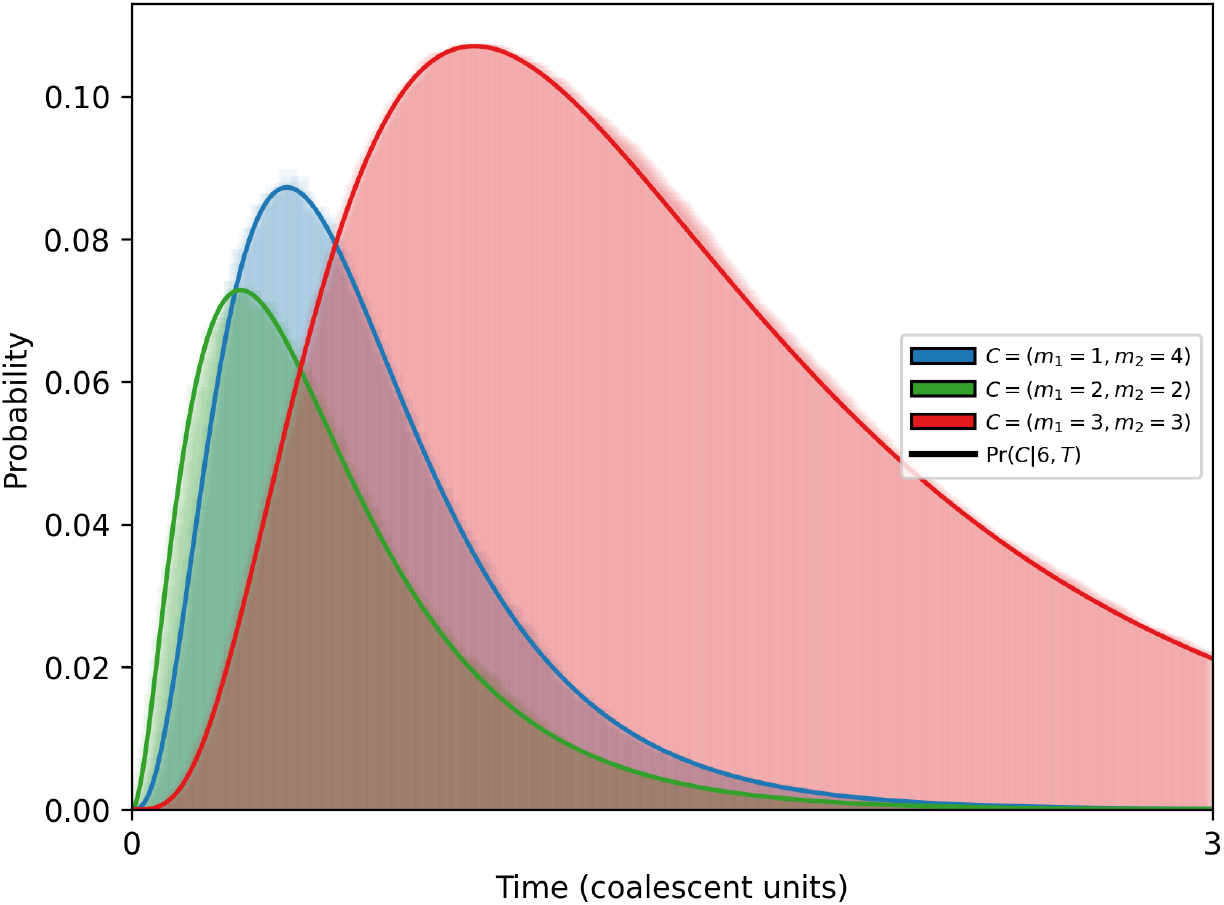
Validating Eq. 5 with simulated data. Here, we simulate the coalescent process and corresponding clade distribution for an example of *n* = 6 sampled lineages. For each time of 500-time points between *τ* = 0 and *τ* = 3, we calculate the probability that *k* randomly chosen clade have sizes *m*_1_, …, *m*_*k*_ as shown in the histogram. Solid curves give analytical expectation from Eq. 5.

**Figure E.9:**
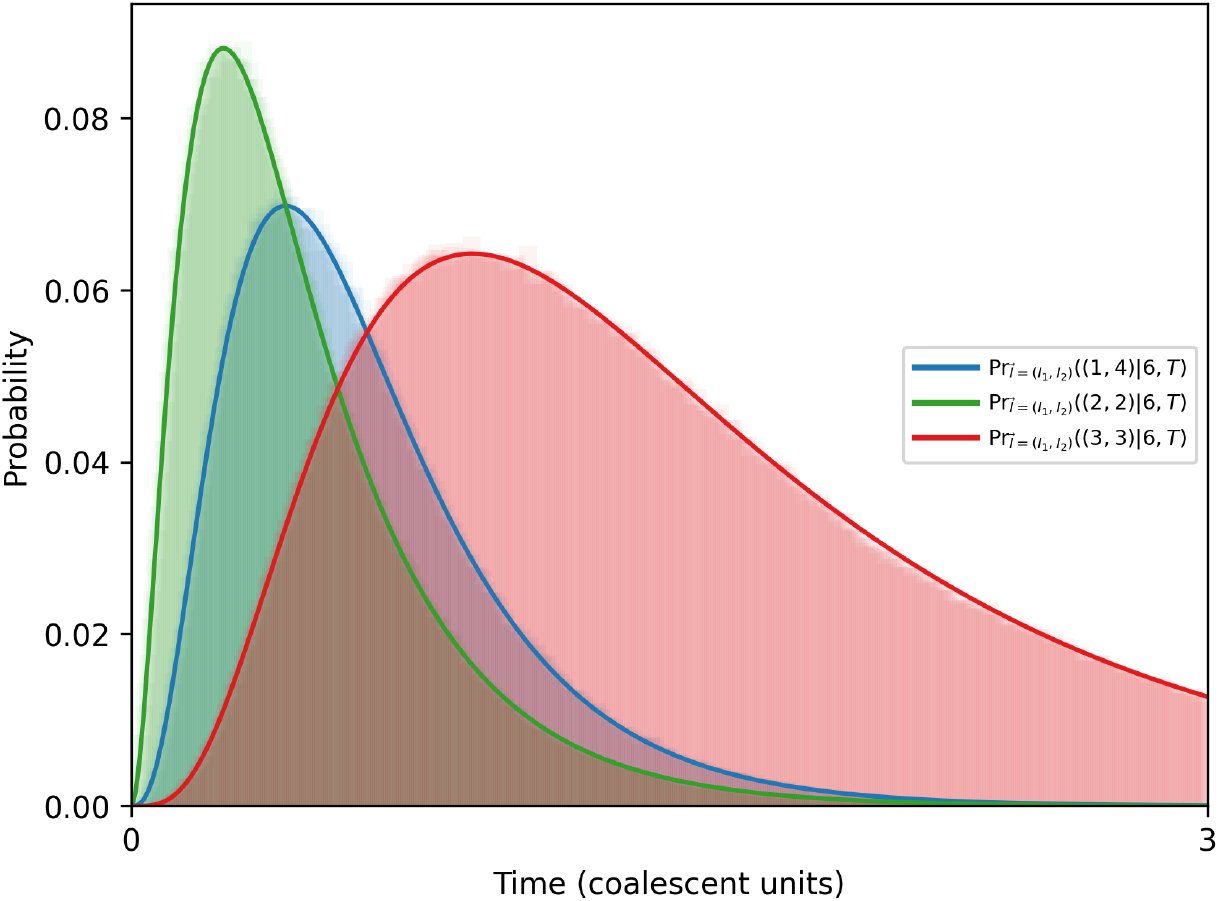
Validating Eq. 7 with simulated data. Here, we simulate the coalescent process and corresponding clade distribution for an example of *n* = 6 sampled lineages. For each time of 500-time points between *τ* = 0 and *τ* = 3, we calculate the probability that *l* randomly chosen individual samples belong to a clade of sizes *m*_1_, …, *m*_*k*_ as shown in the histogram. Solid curves give analytical expectation from Eq. 7.

**Figure E.10:**
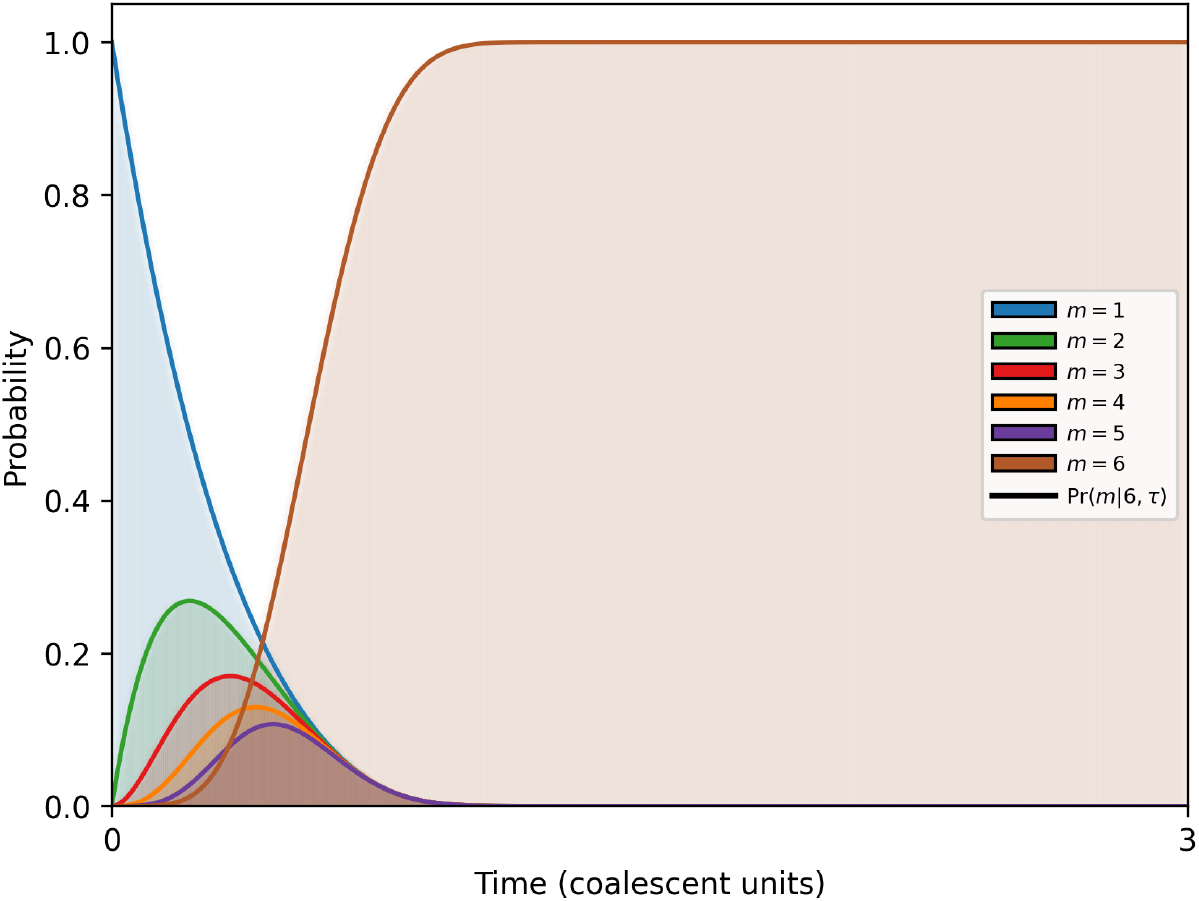
Validating Eq. 9 with simulated data. Here, we simulate the coalescent process and corresponding clade distribution for an example of *n* = 6 sampled lineages. We assume the niche size is growing exponentially at a rate of *r* = 3 and the reference size is *N* = *N*_0_. Solid curves give analytical expectation from Eq. 9.

**Figure E.11:**
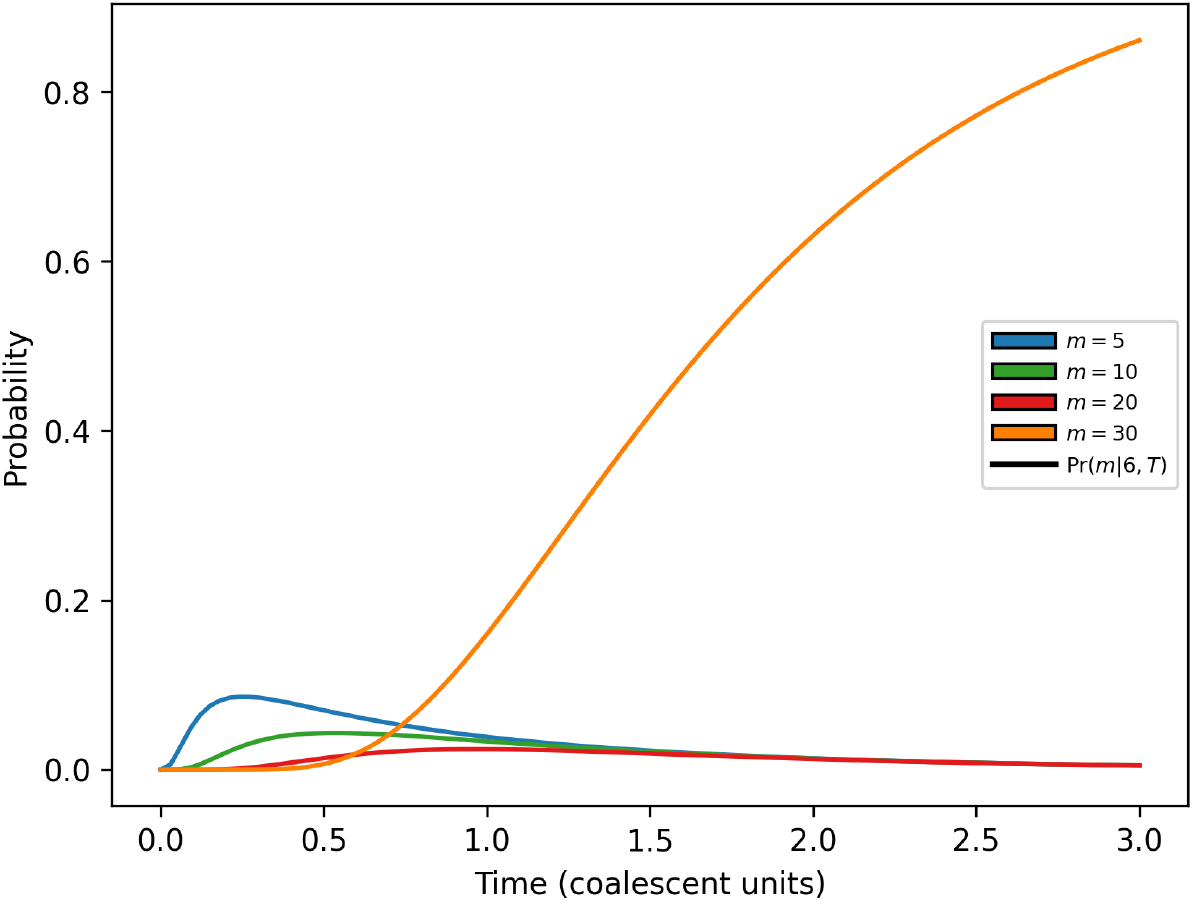
Clade distribution for an example with *n* = 30. We plot the probability of observing 5 clade sizes (*m* = (1, 5, 10, 20, 30)).

**Figure E.12:**
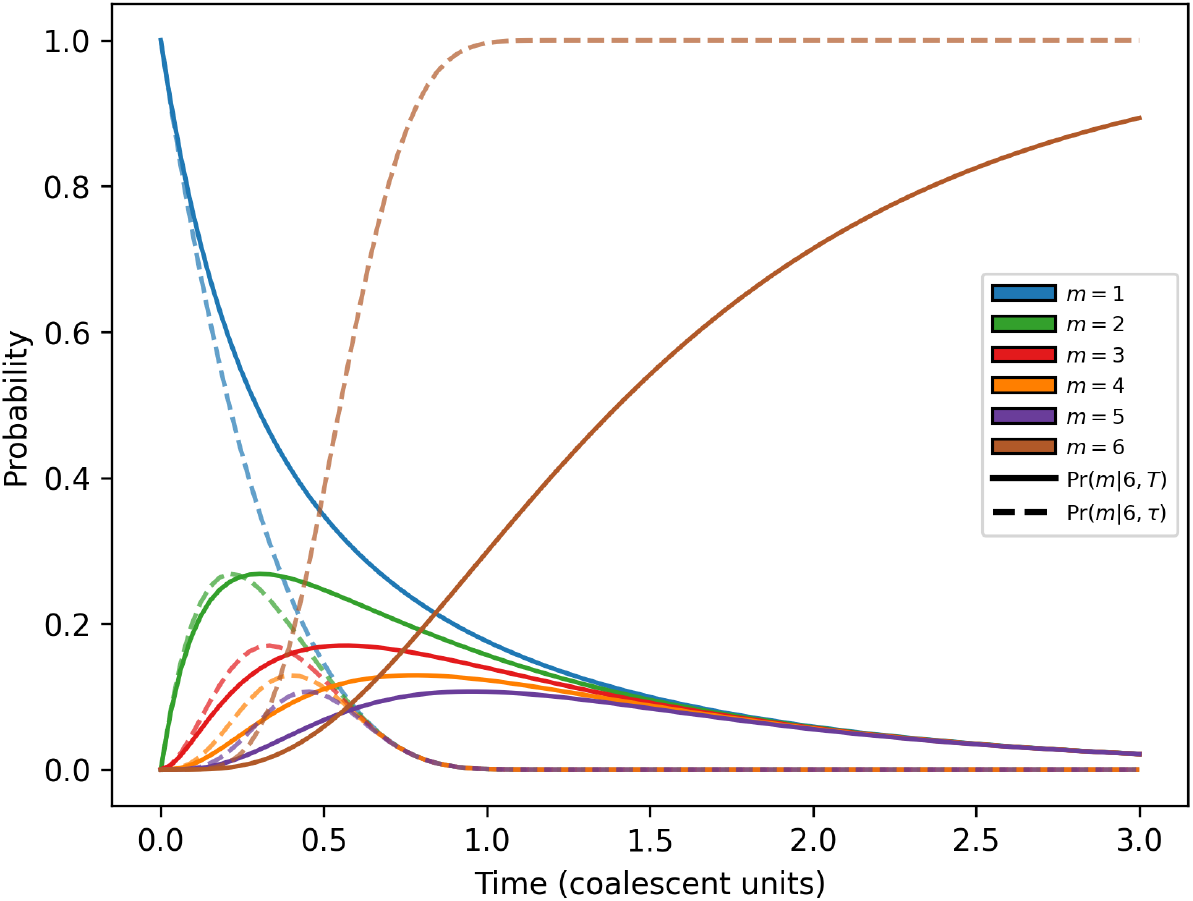
Comparison of clade size in an exponentially growing versus a constant niche size, with reference size at the present day. The distribution of clade sizes over time in a constant niche size *N* (solid) versus in an exponentially growing niche size as given by Eq. 3 and 9, respectively.

